# The level of synovial human VEGFA, IL-8 and MIP-1α correlate with truncation of lubricin glycans in osteoarthritis

**DOI:** 10.1101/2021.03.11.434779

**Authors:** Shan Huang, Kristina A. Thomsson, Chunsheng Jin, Henrik Ryberg, Nabangshu Das, André Struglics, Ola Rolfson, Lena I Björkman, Thomas Eisler, Tannin A. Schmidt, Gregory D. Jay, Roman Krawetz, Niclas G. Karlsson

**Author notes:** Corresponding author - Niclas G. Karlsson, Department of Medical Biochemistry and Cell Biology, Institute of Biomedicine, Sahlgrenska Academy, University of Gothenburg, Sweden. Medicinaregatan 9A, 405 30 Gothenburg, Sweden.

## Abstract

Osteoarthrithis (OA) is an endemic disease due to the increase of the world’s elderly population. Previously thought to be a consequence of an imbalance between cartilage degradation and biosynthesis, it is now recognized as a disease also involving inflammation, hence influencing the level of inflammatory cytokines, growth factors and chemokines. Lubricin is a mucin type molecule where its OA induced glycosylation truncation propels a deteriorating lubrication of the articular cartilage. The objective of this study was to explore the OA driven truncation of *O*-linked glycosylation of synovial lubricin and its cross talk with systemic and local (synovial fluid, SF) inflammation. We compared the systemic level of cytokines/chemokine in OA patients’ and controls’ plasma with their local level in SF using a 44 plex screen. The level of 27 cytokines and chemokines was consistently measured in both plasma and SF. The data showed that the levels of cytokines and chemokines in OA plasma display limited correlation to their counterpart in SF. The level of synovial IL-8 and MIP-1α and VEGFA in OA patients, but not their plasma level, where the only cytokines that displayed a significant correlation to the observed lubricin *O*-linked glycosylation truncation. These cytokines were also shown to be upregulated exposing fibroblast like synoviocytes from healthy and OA patients to recombinant lubricin with truncated glycans mainly consisting of Tn-antigens, while lubricin with sialylated and non-sialylated T anigens did not have any effect. The data suggest that truncated glycans of lubricin, as found in OA, promotes the synovial cytokine production and exerebate the local synovial inflammation.

## Introduction

Osteoarthritis (OA) is a gradual process of cartilage destruction and re-modelling that in combination with synovitis are affecting all areas of synovial joint(1). Since the age of the population is increasing worldwide, OA is an escalating health issue. World Health organization estimate already now that 9.6% of all men and 18.0% of all women aged over 60 years worldwide have symptomatic OA, and predictions indicate that in year 2050, 130 million people will be affected by OA(2). The solution to ease the impact of this prophecy lies in accumulating scientific knowledge of the aetiology of the disease and the events triggering the downwards spiral leading to the chronic destruction of the cartilage. With this knowledge it would be possible to rationalize development of novel treatment required for OA. In this report we are investigating alteration of glycosylation of a protein responsible for lubrication of the joint, lubricin, and the connection between OA related inflammatory markers. Loss of synovial lubrication is one of physiological changes identified in OA and other joint degrading diseases(3).

To date OA has been recognized as a disease involving inflammation with in periods of elevated levels of cytokines/chemokines in the OA joints (4), which can be expressed by chondrocytes, synovial cells, and infiltrated immune cells. These cytokines/chemokines boost joint destruction and activate innervating nociceptors (5). In OA, inflammation of the microenviroment of the joint is believed to be due to decreased blood flow and poor vascularization. This low-grade inflammation is increased with aging and the term “inflammageing” was introduced in the year 2000(6) to explain phenomenon that connects inflammation with aging and associated diseases. The consequence of inflamm-ageing in OA is that chondrocytes experiencing hypoxia and respond by production of reactive oxygen species, proteolytic enzymes and in particular inflammatory cytokines. The proinflammatory cytokine IL-1β is believed to be one of the initiators of OA, inducing downstream synthesis of additional cytokines and regulatory biomolecules(7). Cytokines/chemokines have the potential of triggering signaling pathways that regulate expression of both intracellular and extracellular proteins.

Characteristic for OA inflammageing is this defect in production of extracellular matrix proteins secreted by the classical secretory pathway via the Golgi apparatus glycosylation machinery. That the glycosylation machinery affected is obvious, since shortening of glycosaminoglycans have been detected in OA patients and altered ratio of keratan sulfate/chondroitin sulfate has been reported(8, 9). However, changes of the Golgi are likely to manifest also on other glycoconjugates but these are less investigated. The biosynthesis of the sugar chains is mainly controlled by the activity of their biosynthetic enzymes: the glycosyltransferases. With inflammageing, stimuli such as cytokines/chemokines are altering the expression and location of glycosyltransferases affecting the glycosylation, hence a plausible connection between inflammation and altered glycosylation exists. Disease induced glycosylation is a common theme in glycobiology, where it is most frequently reported in cancer(10). *O*-linked glycosylation alterations in cancer manifest as upregulation of Sialyl Tn-/Tn-antigens and Lewis type epitopes(10). *O*-linked glycosylation changes due to inflammation are less frequently reported; in ulcerative colitis, sialyl Tn expression is upregulated(11) and pulmonary inflammation is associated with decreased expression of blood group antigens, altered sulfation, a switch of core types and increased expression of sialyl Lewis x(12). We have identified that *O*-linked glycans in OA become less sialylated and are more truncated with increased level of T- and Tn-antigens(13).

Lubricin is a higly glycosylated mucinous type molecule, in the synovial fluid encoded by the Proteoglycan 4 (PRG4) gene. Its role is to maintain cartilage integrity by reducing friction at the cartilage surface(14). In addition, lubricin has also been suggested to have growth-regulating properties(15), preventing cell adhesion, and is providing chondroprotection(16-18). Recently, it has also been shown to participate subchondral bone maturation(19). Synovial lubricin is secreted by superficial zone chondrocytes(20), but also originates from synovial fibroblasts(21) and stromal cells from peri-articular adipose tissues(22). Lubricin is also produced by other cells located in tendons, kidney, skeletal muscles, and the ocular surface(23-25). Lubricin has also been shown to be present in plasma(26). The relevance of lubricin in connection to inflammation was highlighted in a mouse model for sepsis, where it was shown to be upregulated in the liver(27), also suggesting that lubricin in plasma is mainly produced by hepatocytes. Here, we investigate the level of inflammation in OA, both systemically in plasma and locally in synovial fluid from OA patients in order to trace the origin of inflammation that is cross-talking with the altered *O*-linked glycosylation found in OA. The investigation was performed using a multiplex cytokine/chemokine screen, in order to identify relevant cytokines/chemokines that are consistently elevated in plasma or synovial fluid (SF) suggesting that they contribute to the OA inflammageing and its association with the altered SF *O*-linked glycosylation. In addition, we investigated if the level of any of the identified cytokines/chemokines that coincided with altered glycosylation of synovial lubricin would be upregulated treating Fibroblast Like Synoviocytes (FLS) with recombinant lubricin (RHPRG4) with truncated glycans. This would indicate a pathway for the cross-talk between inflammageing and altered lubricin SF glycosylation in OA.

## Results

### The cytokine and chemokine profiles in OA SF is different from plasma, both compared to OA patients and controls

SF and plasma were collected from idiopathic late stage OA patients planned for total knee replacemen. We were interested in the inflammatory status and to document the level of inflammatory cytokines/chemokines of these patients. Increased concentration of cytokines and chemokines in our OA patients’ plasma compared to SF would indicate that in addition to the local knee inflammation (measured in SF), the patients would suffer a systemic inflammation (plasma) that could be indicative for OA. In the 44-plex assay we could consistently detect 29 of the components to be present in plasma and among them 27 were presented in SF after our exclusion criteria (Figure 1 and Supplementary Table 1 and 2). Only 8 displayed p<0.05 for altered level between OA and control plasma, 6 with increased and 2 with decreased level in OA plasma. OA and control plasma display similar levels of cytokines/chemokines can be seen in Figure 1, while the levels of cytokines/chemokines in OA SF differed both compared to OA and control plasma (Figure 1). The level of cytokines and chemokines in OA SF mostly displayed increased or similar level as was found in the plasma. The most outstanding exception to this was IL-27 that was found to be higher in both control and OA plasma compared to OA SF.

**Table 1.**
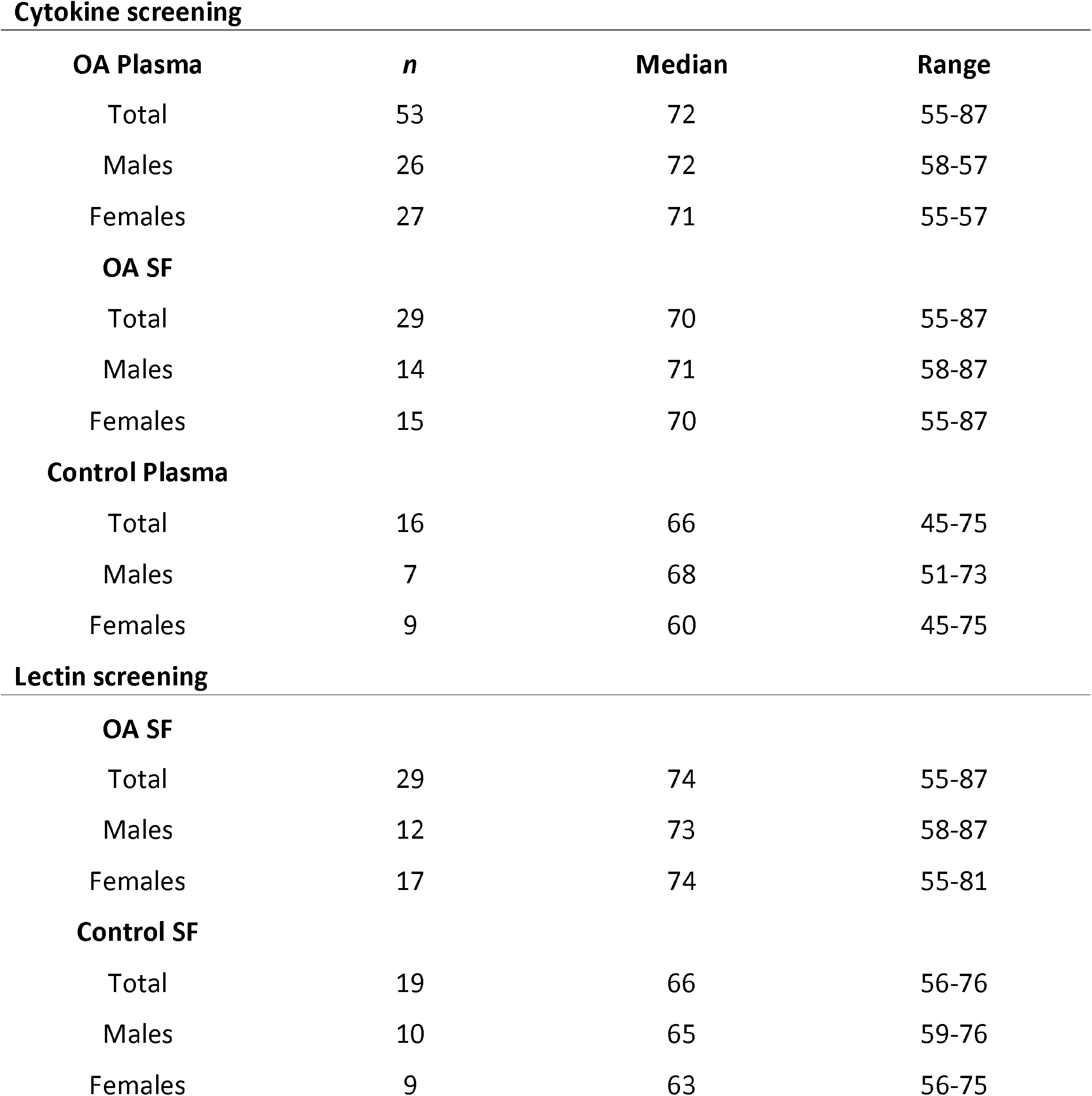
Age and gender distributions of samples collected from OA and controls in the study.

**Cytokine screening**
| **OA Plasma** | ***n*** | **Median** | **Range** |
| --- | --- | --- | --- |
| Total | 53 | 72 | 55-87 |
| Males | 26 | 72 | 58-87 |
| Females | 27 | 71 | 55-57 |
| **OA SF** |  |  |  |
| Total | 29 | 70 | 55-87 |
| Males | 14 | 71 | 58-87 |
| Females | 15 | 70 | 55-87 |
| **Control Plasma** |  |  |  |
| Total | 16 | 66 | 45-75 |
| Males | 7 | 68 | 51-73 |
| Females | 9 | 60 | 45-75 |
| **Lectin screening** |  |  |  |
| **OA SF** |  |  |  |
| Total | 29 | 74 | 55-87 |
| Males | 12 | 73 | 58-87 |
| Females | 17 | 74 | 55-81 |
| **Control SF** |  |  |  |
| Total | 19 | 66 | 56-76 |
| Males | 10 | 65 | 59-76 |
| Females | 9 | 63 | 56-75 |

**Figure 1.**
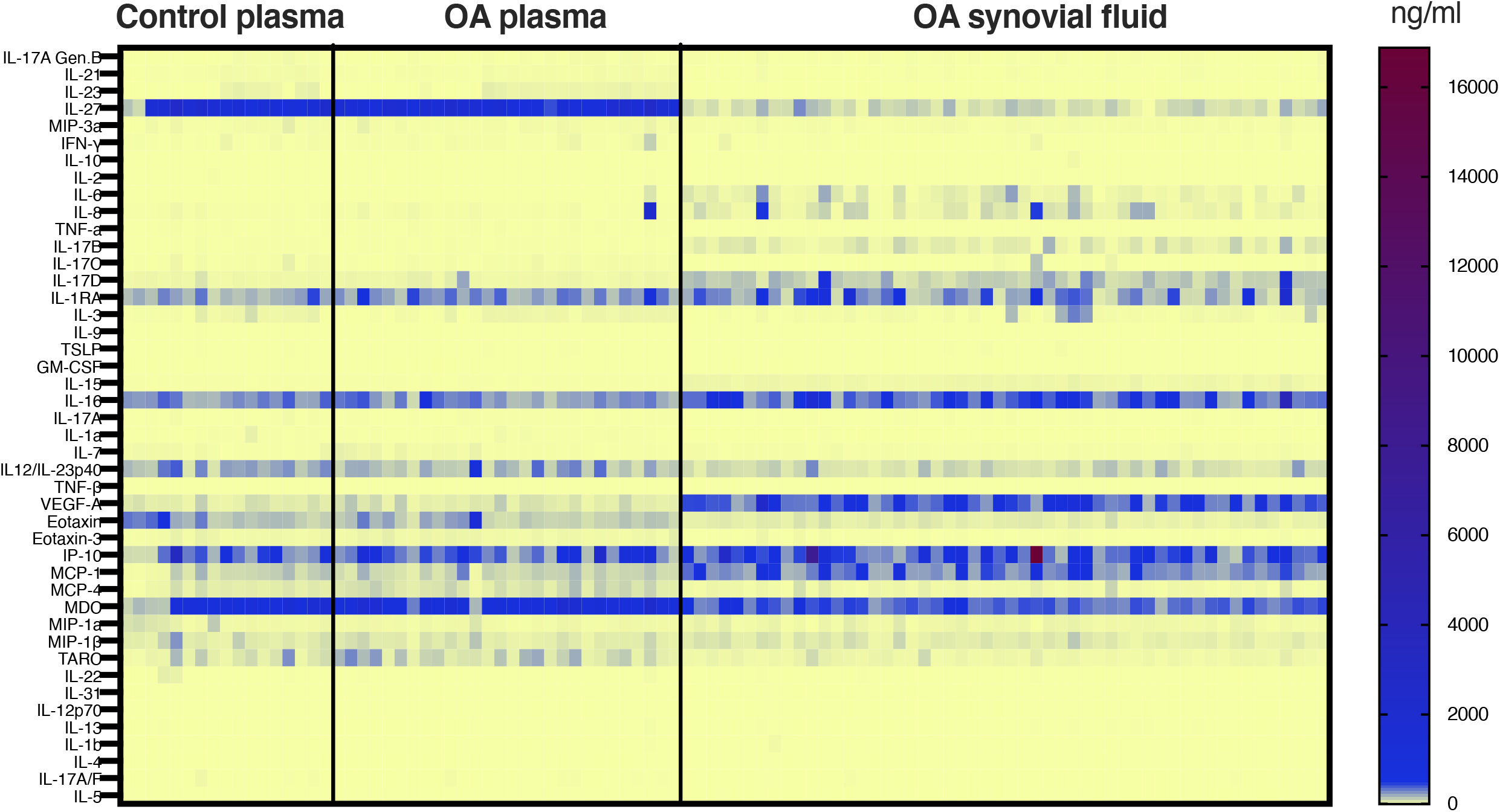
Concentration heatmap of inflammatory cytokines/chemokines expressions in SF and plasma. Analysis was performed in 53 OA synovial fluid, 29 OA plasma, and 16 healthy control plasma. Each row stands for one inflammatory cytokine/chemokine and each column represents each individual (patient/control), and concentration levels (pg/mL) were indicated in color scale.

### MCP-4 and TARC are increased in OA plasma

Comparing plasma from our OA-patient cohort with age-matched healthy control we found that TARC, MCP-4, IL-1RA, MDC, Eotaxin-3 and IL-8 levels were increased in the OA-group (Figure 2, Supplementary Table 1). In addition, the level of Eotaxin and MIP-1α were lower in the OA plasma compared with control. We could also detect a trend of increase for IL-6 (p=0.0688), and IP-10 (p=0.0855) for the OA group. Multiple comparison testing of the differences between OA and control plasma showed that only OA plasma MCP-4 and TARC was significantly increased.

**Figure 2.**
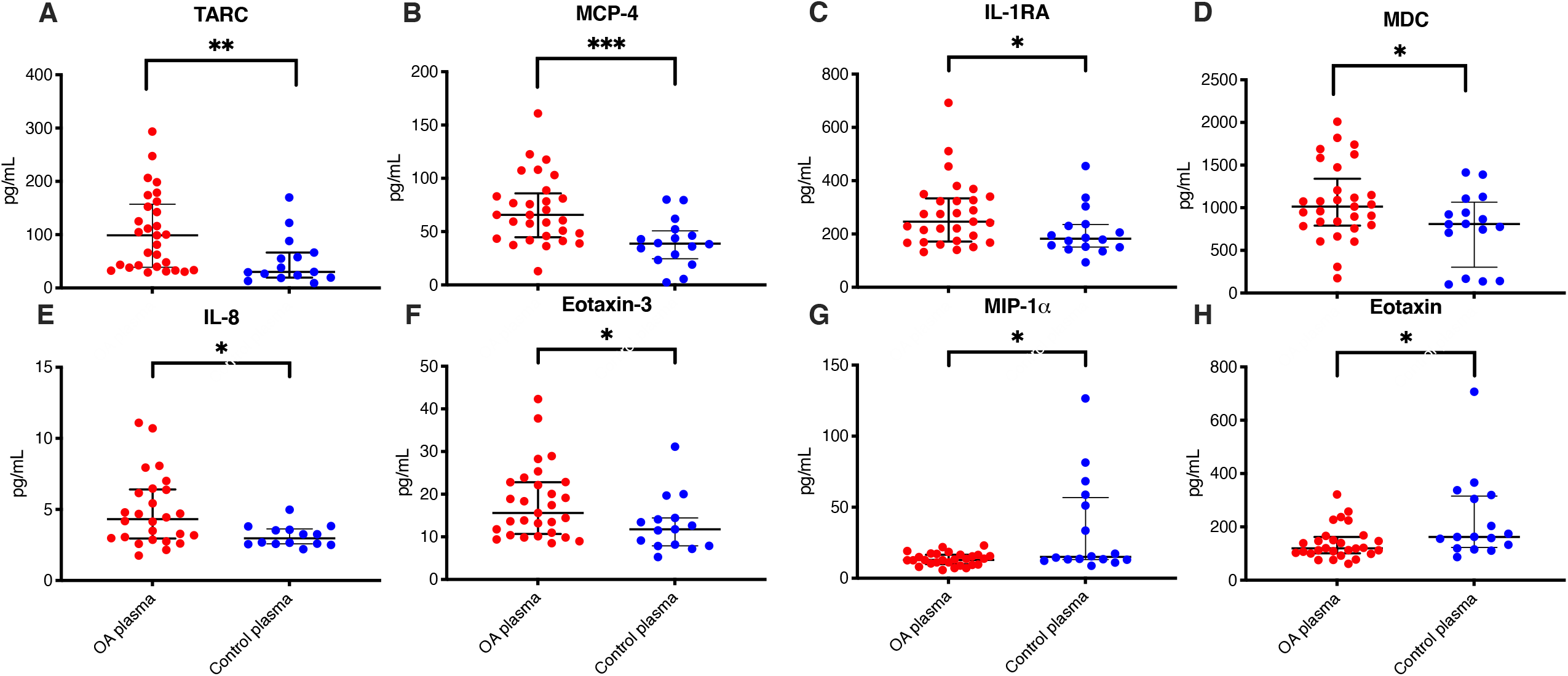
Inflammatory cytokines/chemokines increased in OA plasma versus control plasma. Comparison of the inflammatory cytokines/chemokines level in 29 OA patients’ plasma (red dots) and 16 age-matched healthy controls (blue dots). Data was presented as median with interquartile, and definitive outlier was removed according to ROUT with Q=0.1%.* indicate p <0.05, ** p<0.01, *** p<0.001 and ****<0.0001.

### Comparison of OA SF cytokines and chemokines with plasma and control SF

In order to identify pathological alteration of OA cytokines/chemokines, we compared their levels in OA SF with control plasma and OA plasma and also a with limited number of control SF. Except for IL-21 and IL-1α, the cytokines/chemokines consistently detected in OA plasma were also observed in SF from OA patients. Among these cytokines/chemokines, nine (IL-2, IL-6, IL-8, IL-15, IL-16, IL-17B, IL-17D, MCP-1, and VEGFA) were found with consistent higher level in OA SF compared to OA plasma (Figure 3, Supplementary Table 1). These cytokines/chemokines in OA SF were also significantly higher compared to control plasma. Comparing OA SF with control plasma, only IP-10 was found in addition to the other nine to be increased in OA SF. However, Multiple comparison deemed this observation as not significant. Of the nine cytokines, only IL-8 showed increased level in OA SF compared to OA plasma, and was indicated to be increased in OA plasma compared to control plasma (p=0.0179, but failing multiple comparison stringency) (Figure 2 and 3).

**Figure 3.**
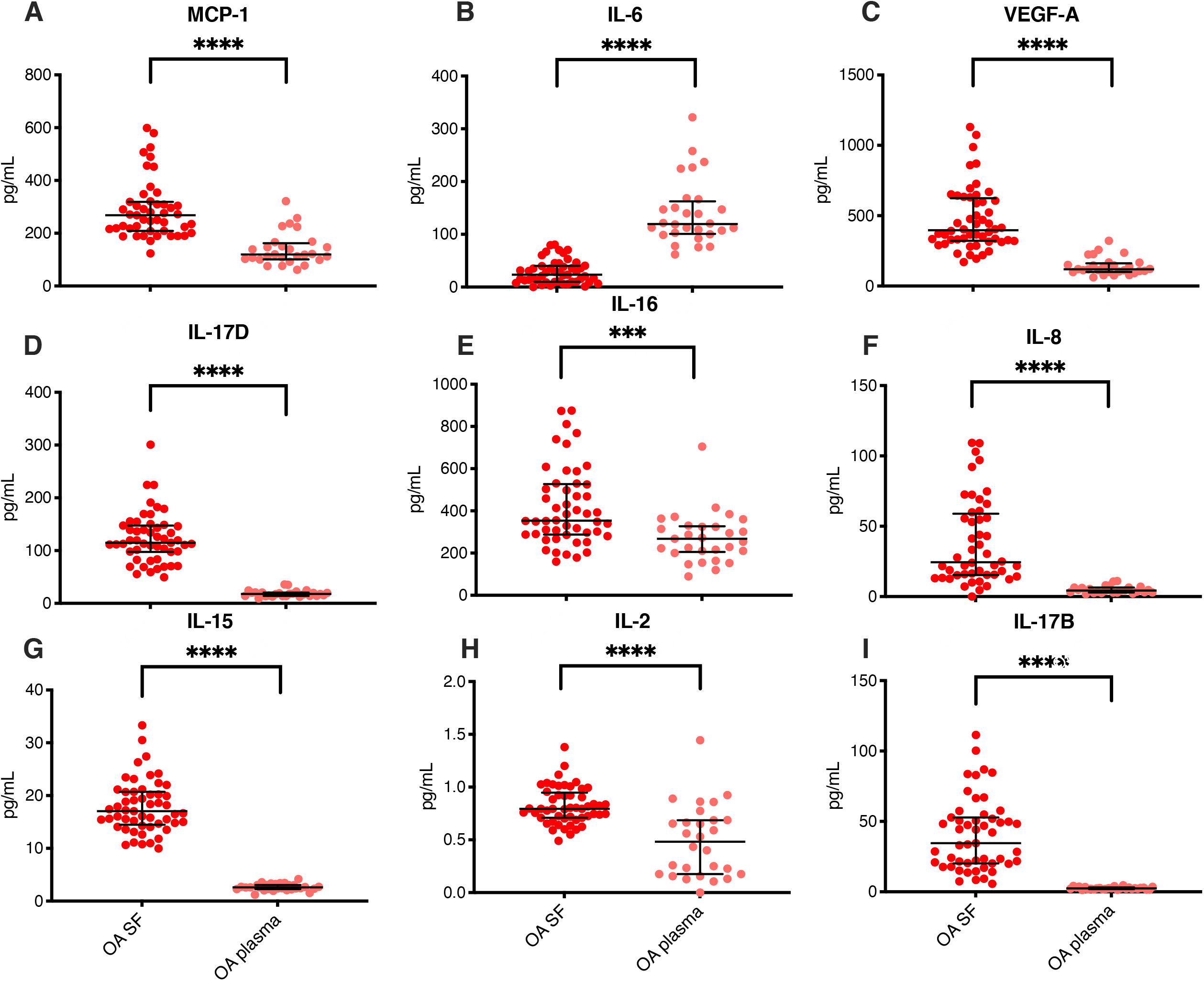
Cytokines and chemokines with increased level in OA SF versus OA plasma. Comparison of the inflammatory cytokines /chemokine level in 53 OA patients’ SF (red dots) and 29 OA patients’plasma (pink dots). Data was presented as median with interquartile, and definitive outlier was removed according to ROUT with Q=0.1%. * indicate p <0.05, ** p<0.01, *** p<0.001 and ****<0.0001.

The data also showed also that the level of TNF-α, IL-3, IL-12/IL-23p40, IL-27, Eotaxin, MDC, MCP-4, Eotaxin-3 and TARC concentrations were significantly lower in OA SF compared to OA plasma (Figure 4 and Supplementary Table 1). MIP-3α was found with a p<0.05 but not fullfilling 5% FDR. Of the significant cytokines, the level of Eotaxin was indicated to be decreased in OA plasma compared to control (p=0.013, but not 5% FDR), while MDC (p=0.0497), MCP-4 (p=0.0010) and TARC (p=0.0030) instead had higher levels (Figure 2 and 4. However only MCP-4 and TARC was found below 5% FDR.

**Figure 4.**
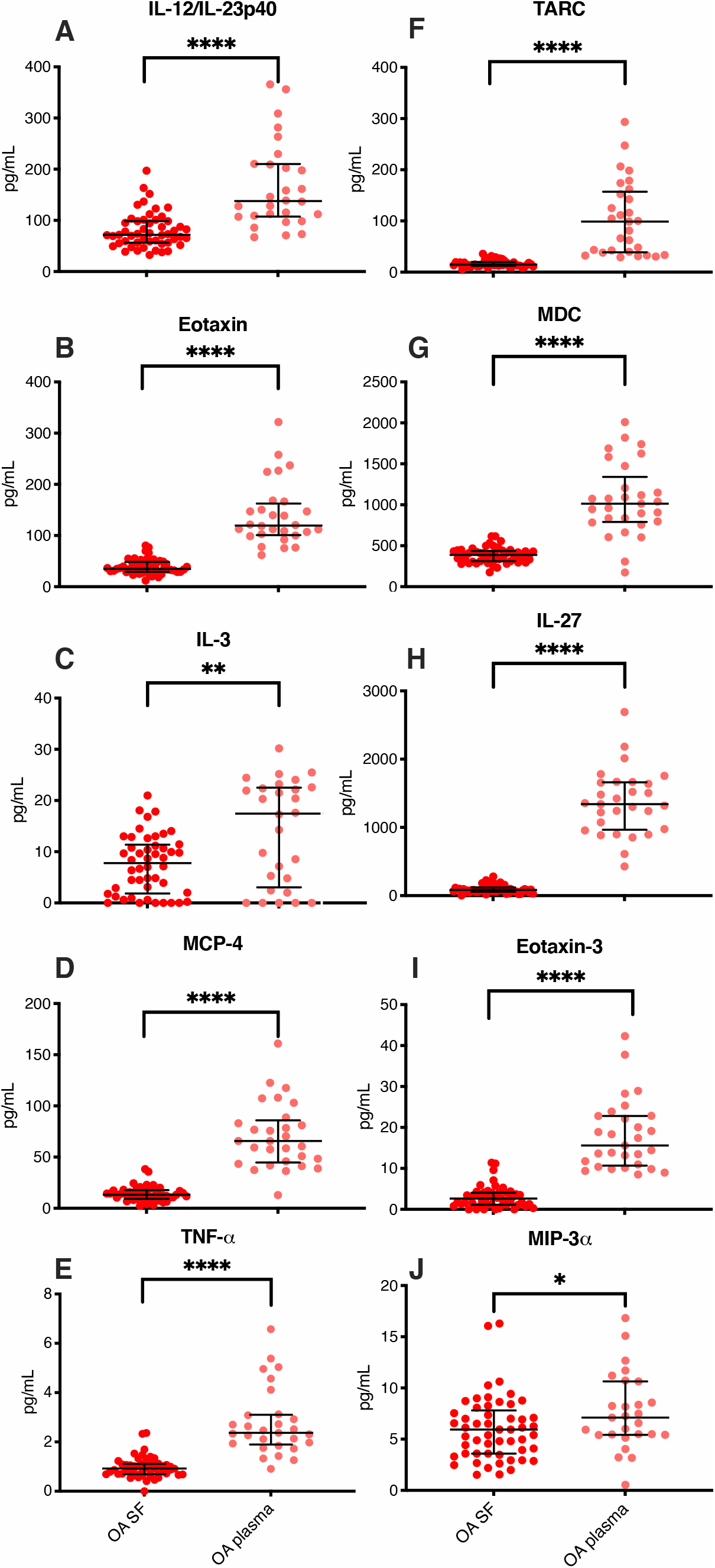
Cytokines and chemokines with decreased level in OA SF versus OA plasma. Comparison of inflammatory cytokine /chemokine levels in OA-SF samples (n=53) (red dots) with OA plasma samples (n=29) (pink dots). Data was presented as median with interquartile, and definitive outlier was removed according to ROUT with Q=0.1%. * indicate p <0.05, ** p<0.01, *** p<0.001 and ****<0.0001.

We also performed a full cytokines/chemokines screen for SF samples from 3 controls (Supplementary Table 2). Several of the cytokines/chemokines that was found to be altered in OA SF compared to control plasma was also altered in the same direction when OA SF was compared to control SF samples. These include IL-6, IL-17D, IL-12/IL-23p40, IL-3 and MCP-1 that would be chategorized as cytokines/chemokines that have little affect systemically when they are altered locally in OA in the joint area, since the levels in OA plasma and control plasma were statistically the same (p<0.05 but rejected after multiple comparison)(Figure 2). We also detected increase of two cytokines, MDC and IL-8, in OA SF compared to control SF, that were also found with an increased level comparing OA plasma with control plasma (p<0.05, but not significant after multiple comparison). MDC was found with more than 10 times higher level in OA SF versus control SF and would be chategorized as a cytokine that is indicated to be increased locally in OA in SF, with only local consequences, since the plasma level (both OA and control) was found to be significantly higher. The other cytokine, IL-8, that was higher in OA SF compared to control SF was found with a 5-10 times higher level in SF compared to plasma (both OA and control) and would be chategorised as a cytokine effecting both locally and systemically. In addition, we detected a decrease of MIP-1β and IL-7 in control SF compared to OA SF (p<0.05). Their level in OA SF were not significantly different compared to OA and control plasma, and not significantly differnt between OA and control plasma. Again, this indicated that a local alteration in SF cytokines/chemokines level may not influence the plasma level.

### Correlation of cytokines/chemokines within and between plasma and SF

We further investigated how inflammatory cytokines/chemokines were co-regulated in the same fluids (SF or plasma). Twenty-seven expressed cytokines/chemokines in all test fluids (control plasma, OA SF, OA plasma) were subjected to the correlation test (Supplementary Figure 2). Overall, we found that the level of the measured cytokines/chemokines were positively correlated, with significant positive value of the average Spearman correlation in OA SF, OA plasma and control plasma (Supplementary Figure 1). This is anticipated, since low a low level inflammation in OA and controls will manifest as increased level of multiple inflammatory markers. However, OA SF levels of cytokine/chenokines were found to be less correlated compared to control plasma, but not OA plasma, and no signicant differences was found between OA and SF plasma.

In order to identify potential cytokines/chemokines that were able to diffuse between plasma and SF, correlations between matched SF and plasma cytokines/chemokines level from the same patient levels were plotted and correlation calculated for the 27 detected cytokines/chemokines in all test fluids. Correlations between SF and plasma were identified for TNF-α, IL-12, IL-10, IL-27, IP-10, MDC, and TSLP (Figure 5). However, TSLP and IL-27 was not found to be significant after multiple comparison.

**Figure 5.**
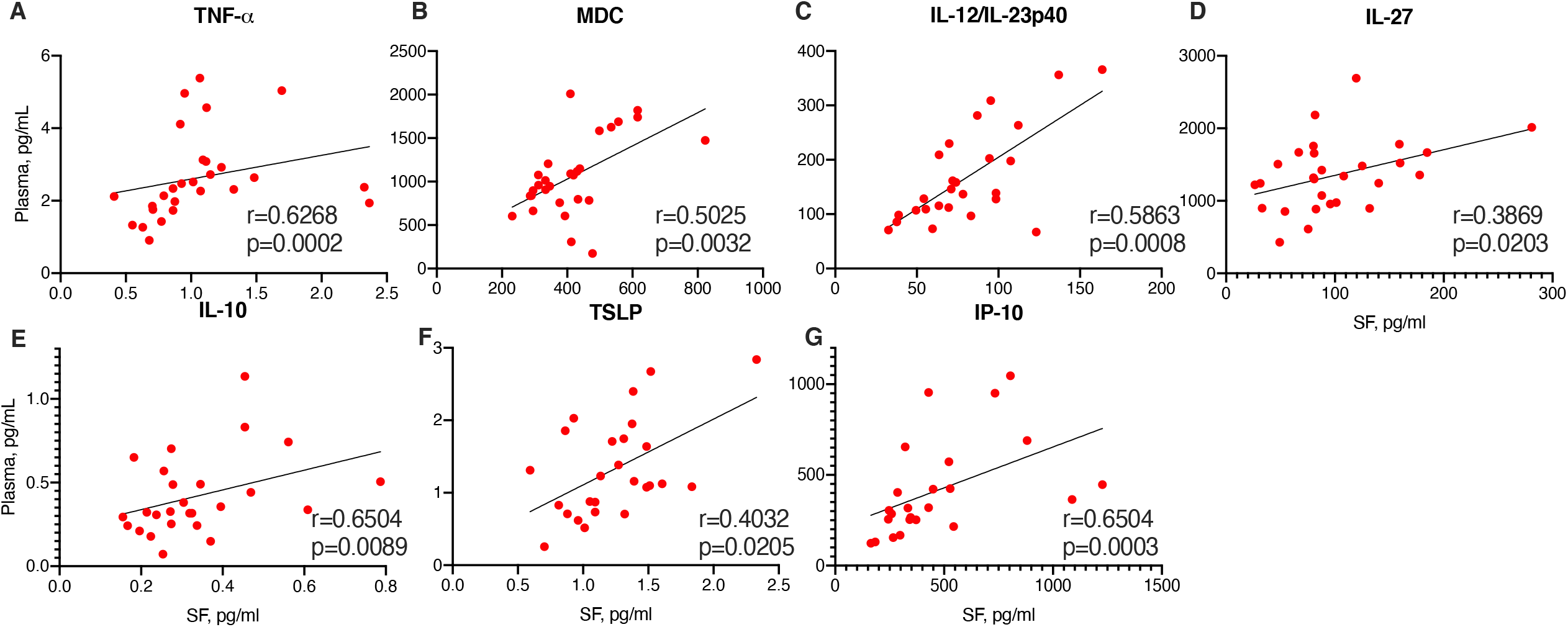
Correlations between cytokines/chemokines expression in synovial fluid and plasma in OA. Correlations between inflammatory cytokine expression in synovial fluid and plasma from 28 OA patients. Spearman correlation r and significance p were calculated and indicated in each diagram. Outliers were removed according to ROUT with Q= 0.1%.

### *O*-linked glycans on SF lubricin displays increased level of Tn- compared to T- antigens in OA

We investigated if the glycosylation of SF lubricin varied between our samples and between or OA and control cohort. First we measured using LC-MS the level of two main oligosaccharides released from OA SF lubricin, GalNAc and the Galβ1-3GalNAc, where an increase of the former would indicate increased truncation of lubricin glycans (Supplementary Figure 2). Secondly, we measured the same epitopes using a lectin/antibody sandwich ELISA, where the glycans where still attached to lubricin.We found that the sandwich ELISA and LC-MS provided similar results (Figure 6A), with a direct correlation (p=0.016) between the two readouts. This allowed us to predict the glycosylation change based on lectin response in the more sensitive ELISA assay in our scarce and low volume control SF samples. Using ELISA on SF from 29 OA and 19 non-OA control we could infer that SF lubricin on our OA patient cohort contained significant more Tn antigens compared to T-antigens (Figure 6B). This showed that the oligosaccharides on OA lubricin are shorter than the controls, in line with previous publication(13).

**Figure 6.**
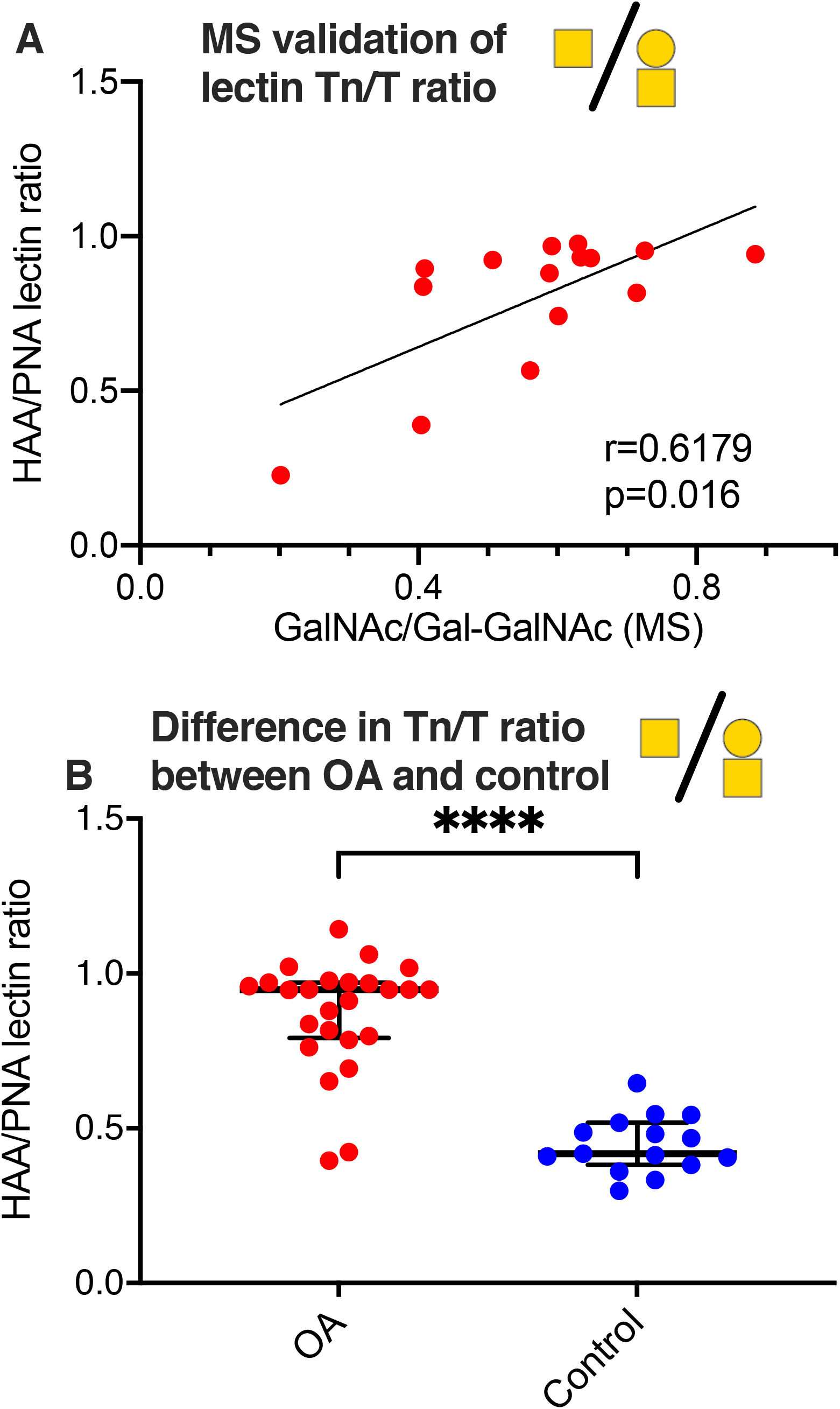
Determination of Tn and T antigens on SF lubricin. (A) Correlation between Tn and T antigens using MS or lectins, where the level of Tn antigen (GalNAcα1-Ser/Thr) where measured using the lectin HAA and the level of T-antigen (Galβ1-3GalNAcα1-Ser/Thr was measured using the PNA lectin using patients diagnosed with OA (n=16). Spearman correlation r and significance p were calculated and displayed in the diagram. (B) Absorbances of HAA-epitopes/PNA-epitopes on SF lubricin were calculated from 29 OA patients and 16 controls using lectin ELISA. Significance were calculated by two-tailed nonparametric Mann-Whitney test. Outliers in were excluded prior calculations according to GOUT test with Q= 0.1. Cartoon of monosaccharide building blocks for representing oligosaccharides according to the SNFG nomenclature, yellow circle=Gal, yellow square=GalNAc.

### Cytokine/chemokine expressions correlated with glycosylation features of lubricin in OA

We further went along and investigated if the observed glycosylation change in OA was associated with the inflammation markers in SF, since our data inicated that systemic inflammation markers has limited influence to the SF inflammation and a potential cross talk with the glycosylation of SF lubricin. We found that HAA/PNA ratio correlated with the inflammatory cytokines/chemokines MIP-1α, MIP-1β, MCP-1, Il-8, VEGFA, and MDC expressions in OA SF (Figure 7A-F) and Supplementary Table), but only IL-8, VEGFA and MIP-1α, was shown to be significant after multiple comparison. Aligned with our findings, the results from non-OA control SF fit with the trends that was observed for OA SF (Figure 7A-F, blue dots), i.e. low expression of these cytokines/chemokines is associated with degreased HAA compared to PNA. This suggest that increased amount of shorter glycans on lubricin is associated with high expression of certain expression modulating mediators in SF with an emphasis on IL-8, VEGFA and MIP-1α.

**Figure 7.**
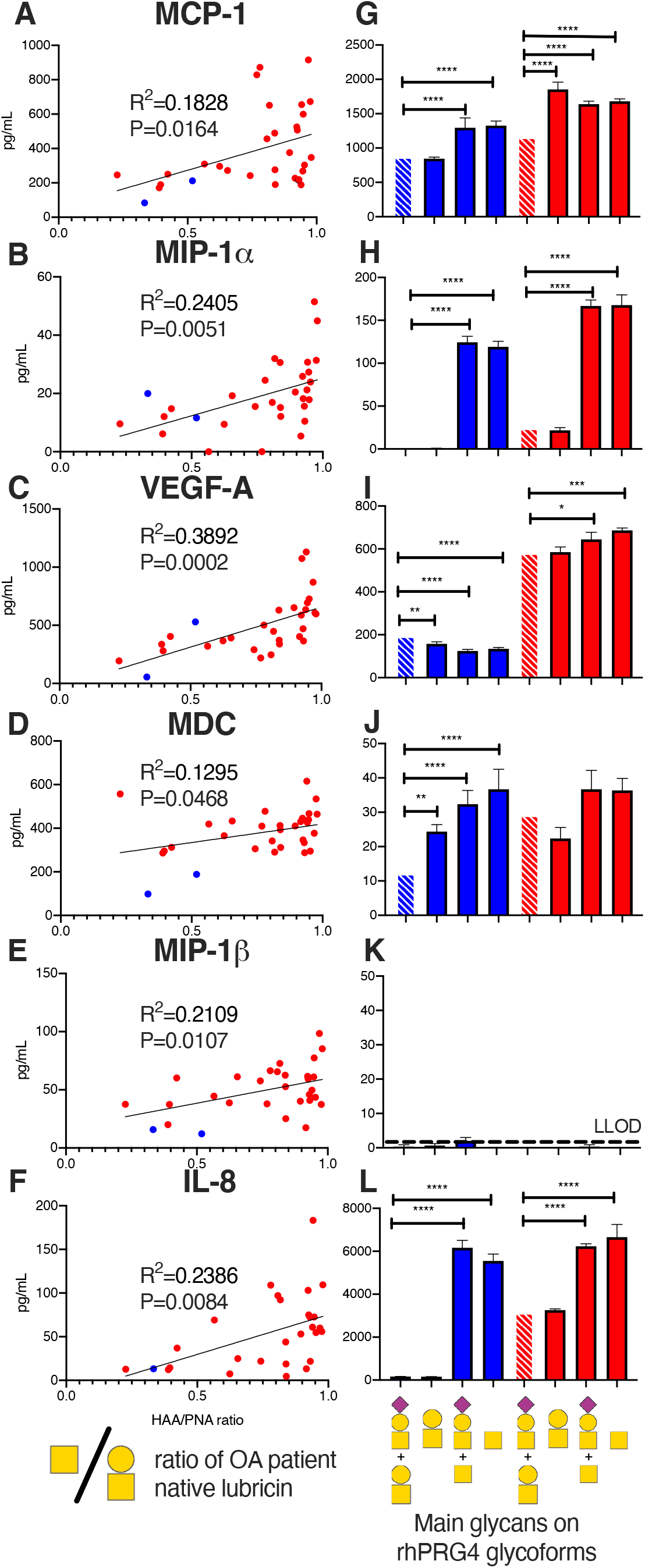
SF lubricin glycan modification in OA increases inflammatory biomarker expression. Left hand panels (A-F) shows correlations between HAA/PNA ratio and inflammatory cytokine expression in the same OA patient (red dots, n=29) including also non-OA controls (blue dots, n=2). ELISA for each sample was measured in duplicates. Outliers in were excluded prior calculations according to GOUT test with Q= 0.1. Spearman correlation r and significance p were calculated and displayed in each diagram. Right hand panels (G-L) shows the cytokine level after treatment of fibroblast like synoviocytes (FLS:s) isolated from healthy individuals (n=3, blue bars) and OA patients (n=3, red bars) using recombinant lubricin (rhPRG4) with or without treatment with β-galactosidase and sialidase. In L is the main oligosaccharides present on rhPRG4 shown with or without digestion corresponding to (from left), untreated in PBS, treated with sialidase, treated with β-galactosidase and treated with both sialidase and β-galactosidase of FLS from controls (blue) and OA patients (red Cartoon of monosaccharide building blocks for representing oligosaccharides according to the SNFG nomenclature, yellow circle=Gal, yellow square=GalNAc, purple diamond=NeuAc. Significance of secretion from FLS was calculated using multiple comparison one-way ANOVA compared to FLS (healthy and OA, striped bars) treated only with recombinant lubricin. * indicate p <0.05, ** p<0.01, *** p<0.001 and ****<0.0001. Secretion of cytokines from FLS using controls subjected to PBS or PBS + enzymes was found not statistically different compared to treating the FLS with intact rhPRG4.

### Desialylated and/or degalactosylated lubricin stimulates cytokine secretion of FLS

Since our data suggested that lubricin glycans became truncated in OA, displaying more Tn-antigens, we set out to test the hypothesis that lubricin with truncated glycans are able to stimulate cytokine expression in cells from the synovial area. We generated glycoforms of lubricin by treating recombinant lubricin containing sialylated core-1 structures(13) using sialidase and/or β-galactosidase. Treating FLS cells isolated from healthy individuals and OA patients with these lubricin glycovariants, we could detect that the cytokines that were identified correlating with increased truncation(MIP-1α, MCP-1, Il-8,VEGFA, and MDC) were effected (Figure 7G-L). The only exception was MIP-1β, where the background secretion was low, and no effect could be detected after treatment. The effect was most pronounced on IL-8 and MIP-1α. For these two the effect appeared to be directly an effect of the appearance of Tn-anigens, since the β-galactosidase with our without sialidase showed a significant increase, while only sialidase did not have any effect.

The effect of treating FLS using of lubricin with truncated glycans had an opposite effect on VEGFA compared to IL-8 and MIP-1α since it was found to decreas its VEGFA expression from an already low lewel in control FLS. In contrast, FLS from OA patients already without treatment showed a high VEGFA secretion, and after increasing the amount of Tn-antigens (using β-galactosidase and/or sialidase) on rhPRG4, the production increased even further. Again this increase was not seen using only sialidase. MDC and MCP-1 that was displaying the lowest correlation with lubricin in patient samples, were also showing a mixed results concerning the increase of secretion due to increase of Tn-antigens. MCP-1 from control FLS and from OA donors was shown to increase due to increase of Tn-antigens. However for OA-donors, just using desialyation (increase of T-anigens) provided the highest effect in increasing the MCP-1 secretion. For MDC, truncation of recombinant lubricin glycans using either of the exoglycosidases individually or together increased the expression in FLS from healthy individuals, while there was no significant change on the secretion for the FLS from OA individuals. The conclusion is that truncated glycans on lubricin are able to sustain a inflammation by triggering specific inflammatory cytokine expression of synovial cells.

## Discussion

### Cytokines and chemokines in knee-OA

The cytokines and chemokines data comparing SF/plasma of OA patients and controls suggests that the level of specific but not necessarily the same cytokines/chemokines where pathologically altered in either SF or plasma or sometimes both (Fig. 2-4). Hence, based on the cytokines and patients analysed here, the specific inflammation detected in OA SF is not reflecting a systemic inflammation measurable as altered level of plasma cytokines/chemokines. The unique cytokine/chemokine environment of the SF is only partly influenced by the cytokine/chemokine environment of plasma. Our data also verifies that a local inflammation persists in the OA joints.

MCP-4 and TARC were the only cytokines/chemokines in this study shown to be significant comparing OA and control plasma in knee-OA. These cytokines/chemokines were not analysed in a previous cytokine screen of OA versus control serum(28). In previous report, differential level of Eotaxin, MCP1, MIP1β and VEGFA were detected, but could not be verified here, although Eotaxin showed a tendency of being increased.

TARC, that was found in this report to be upregulated in OA plasma, is known to induce T cell activation(29), and has been suggested to be linked to the granulocyte macrophage-colony stimulating factor dependent inflammatory pain(30). The other elevated OA plasma chemokine; MCP-4: has previously been found to be elevated in knee-OA serum and associated with the severity of cartilage degradation(31). Both TARC and MCP-4 belong to CC chemokine that stimulate the migration of monocytes and other cell types such as NK cells and dendritic cells. Neither the level of TARC nor MCP-4 have to our knowledge been studied systematically in SF in an OA context. With only a limited number of control SF analyzed here (Supplementary Table 2), the results are inconclusive about the alteration of MCP-4 and TARC in OA SF. However, the plasma level of both are significantly higher compared to OA SF, indicating that the elevated levels of these chemokines in plasma are not due to increased synovial biosynthesis, but are rather representing an increased systemic inflammation in late stage OA.

Strong (both positive and negative) correlations of cytokines and chemokines within each fluid analysed (OA SF and plasma and control plasma) were found between cytokines and chemokines in the control plasma samples, but positive correlation between cytokines/chemokines in OA plasma, control plasma and OA SF dominated (Supplementary Figure 1). Interestingly, we found that IL-17B and IL-17D were consistently negatively correlated or un-correlated with the other cytokines/chemokines measured (Supplementary Figure 1). IL-17B has been shown to be the dominant IL-17 cytokine in synovial neutrophils in rheumatoid arthritis patients(32). However, our data based on the other inflammatory markers is more consistent with that neutrophil driven inflammation is only a minor contributor to OA synovial tissue inflammation. IL-17D has also been shown to be shown to be secreted by synovial neutrophils but to a lesser degree(32). Its positive correlation with IL-17B may suggest that it is of neutrophil origin in a subtype of OA patients with comorbidities, where the inflammation of the joint in mostly neutrophil driven.

To identify potential communication of cytokines/chemokines between SF and plasma we investigated the correlation of cytokine/chemokine levels between plasma and SF (Fig. 6). Only positive significant correlations of the cytokines/chemokines were found between plasma and SF, reassuring its significance. However, since the majority of the 27 cytokines and chemokines displayed significant differences between SF and plasma but no correlation found (Figure 3 and Supplementary Table 1), one interpretation would be that the cytokine/chemokine flux between synovial fluid and plasma is restricted. Alternatively, it is influenced by effects from removal (degradation, absorption, biological barrier), synthesis and/or secretion of cytokines and chemokines effecting the level in plasma and/or SF specifically.

### Cytokine/chemokine and lubricin glycosylation

Our data from cytokine/chemokine environment of the SF suggest that it harbors a unique profile separate from the plasma. Most of the differences cannot be explained by in-flux from plasma. Hence, synovial cells has to be able to regulate its own cytokine/chemokine environment, and as such, crosstalk between glycosylation of lubricin and cytokines/chemokines is regulated localy in the joint. In this paper and in a previous paper(13) we showed that *O*-linked glycans are significantly shortened on synovial lubricin in late stage OA, increasing the number of exposed Tn-antigens. This exposure would allow OA synovial lubricin to establish different type of interaction with synovial cells. Previously it has been found that lubricin can interact with CD44(33), where there was indication that interaction increase with deglycosylation. However, this interaction was belived to be due to an increased exposure of lubricin protein backbone allowing more efficient interaction with CD44, and not an effect of a direct interaction between a synovial receptor and exposed Tn-antigens on lubricin. Toll-like receptors (TLRs) has been found to be present on FLS and fullfills many of the criteria required to be the candidates to be involved for triggering a cytokine release after Tn lubricin stimulation. TLR2, TLR3, TLR4, TLR-5 and TLR9 has been found on FLS isolated from OA and RA patients(34, 35), where the TLR receptor ligand interaction was shown to induce IL-6, IL-8, TNF-α and VEGF(34), and stimulation(34). Stimulation with of TLRs with LPS together with sialylated recombinant lubricin enhanced the MIP-1α, expression, while sialylated lubricin shown previously(35) and here, did not have an effect. The Tn-antigen appears to be a very potent stimuli of innate immune system, where Tn-containing mucins and Tn-antigen coated nano-particles have been shown to trigger IL-6 secretion of monocytes(36, 37) and acting as a co-stimmulator of TLR mediated innate immune respone(37).The ability of FSL to be able to trigger the identified IL-8, MIP-1α and VEGFA secretion by Tn-antigen stimmulation via a pathway involving TLRs has not in our knowledge been investigated.

The question for how and when the lubricin glycosylation is modified during the OA development remains open and may also involve a cytokine/Tn-antigen crosstalk, where cytokines stimulate altered glycosylation of synovial cells, via unknown pathways. The only candiates correlating with glycosylation change was again VEGFA, IL-8 and MIP-1α. VEGFA has been suggested to be associated with core-2 GlcNAc-transferase in ovarian cancer, responsible for synthesizing core-2 and -4 *O*-linked oligosaccharides(38). Core-2 GlcNAc transferase has also been indicated to be increased by TNF-α stimulation of bovine synoviocytes(39) and human chondrocytes(40) in-vitro, the latter also including increase of the core-1 galactosyltransferase. IL-8 together with Il-6 has been shown to induce glycosyltranferase and sulfotransferase expression in bronchial tissue explants in vitro(41). TNF-α as well as IL-6 have been shown to up-regulate Tn levels in gingival fibroblasts, probably by downregulation of the COSMC chaperon required for activation of the core-1 galactosyl transferase that allows *O*-linked glycans to be elongated. This latest example would provide a the type of truncation of the glycosylation profile consistent with what has been observed in on OA lubricin in this report and previous(13). MIP-1α is a chemokine with limited information about its association with OA. It has been reported to be secreted by vast majority of white blood cells(42) but also epithelial cells has been shown to secret MIP-1α after IL-1β stimulation(43). In relation to synovial tissue, synovial fibroblast has been shown to induce MIP-1α after TNFα stimulation in vitro, while IL-8 was not(44).

Both Il-8 and VEGFA that corelated with glycosylation change has the cabability to both alter the glycosylation as well as its secretion is stimulated by glycosylation truncation, points towards this cross-talk between inflammation and glycosylation. However, the ability of MIP-1α to cross talk with glycosylation is unexplored, and if it is capable of also trigger glycosylation change is unknown.

The hypothesis that arise from the data in this report about the connection between cytokines/chemokines and glycosylation crosstalk of lubricin in OA is that initial synovial cytokine/chemokine expression trigger the increase of Tn-antigen expression probably by the by the prime cytokines in OA, i. e. IL-1β and TNF-α. This creates an inflammation downwards spiral in OA, where the Tn-antigen itself propagates the OA local joint inflammation further, by inducing additional pro-inflammatory cytokines.

## Material and methods

### Patients and samples

SF and EDTA plasma samples for cytokine screening and lectin screening were collected from late-stage OA patients that were subjected to knee replacement surgery (patient informationn in Table 1). The samples were collected prior surgery, centrifuged, aliquoted, and stored at -80°C until assayed. Control plasma was collected from age matched individuals without reported history of OA, arthritis or reported trauma of joints. Control SF were aspirated from cadaveric donations without history of arthritis, joint injury or surgery, no prescription anti-inflammation medications, no co-morbidities, and were collected within 4 hours of death. Inclusion criteria for control cadaveric donations for collection of normal group fibroblast-like synoviocytes (FLS) (n=3): age of 18 years or older, no history of arthritis, joint injury or surgery (including visual inspection of the cartilage surfaces during recovery), no prescription anti-inflammatory medications, no co-morbidities (such as diabetes/cancer), and availability within 4 hours of death. Inclusion criteria for knee OA (n=3) was based on a diagnosis of OA performed by an orthopedic surgeon at the University of Calgary based on clinical symptoms with radiographic evidence of changes associated with OA in accordance with American College of Rheumatology (ACR) criteria. Radiographic evidence of OA was collapsed or near collapsed joint space of any compartment of the knee. All samples were collected after written consent from patients and controls and the procedure was approved by the Conjoint Health Research Ethics Board of the University of Calgary (REB15-0005 and REB15-0880) or by the Regional Ethical Review Board in Gothenburg (ethical application 172-15) and the study abide by the Declaration of Helsinki principles.

### Cytokines/chemokines multiplex ELISA validation in synovial fluid samples

Concentrations of inflammatory cytokines/chemokines in SF and plasma were measured by Meso Scale Discovery (MSD) V-PLEX Human Cytokine 44-Plex immunoassay kit (Gaithersburg, MD, US) according to the manufacturer’s instructions. SF Assay validation was applied by using two SF samples to measure linearity of dilution and spiking. The synovial fluid samples were diluted 1:2 into 5 different dilutions in sample diluent provided by manufacturer) and loaded in duplicates to test for linearity of dilution. Expected concentrations were calculated as a mean concentration of 5 preparations and recovery rate was calculated for each sample preparations. Spiking validation was carried out by adding 3 calibrators with known concentrations (recombinant standard for each cytokine/chemokine that were diluted for high, medium, and low concentrations with standard diluent, provided by manufacturer) to the tested two synovial fluid samples. Expected values were calculated by calibrator concentration plus expected synovial fluid concentration from the dilution linearity test, and recovery rate was compared between experimentally read and calculated (Supplementary Data). For Limit of Quantification (LOQ), the manufacturers reported limits were used rangeing between 0.45-713 pg/mL with an average of 23.4pg/mL and with a typical inter and intra plate variation of <10%.

### Analysis of regulatory markers in OA and control

Twenty-nine OA plasma, 16 healthy control plasma, 53 OA SF and 3 non-OA control SF samples were measured in 5 multiplex panels in dublicates using the V-PLEX Human Cytokine 44-Plex (MSD, Rockville, MD, USA). Concentrations of following inflammation cytokines/chemokines were measured in each sample: Eotaxin, Eotaxin-3, Granulocyte-macrophage colony-stimulating factor (GM-CSF), interferon (IFN)-γ, interleukin (IL)-10, IL-12/IL-23p40, IL-12p70, IL-13, IL-15, IL-16, IL-17A, IL-17A/F, IL-17B, IL-17C, IL-17D, IL-1 receptor antagonist (RA), IL-1α, IL-1β, IL-2, IL-21, IL-22, IL-23, IL-27, IL-3, IL-31, IL-4, IL-5, IL-6, IL-7, IL-8, IL-9, Interferon gamma-induced protein (IP)-10, monocyte chemoattractant protein (MCP)-1, MCP-4, Macrophage-derived chemokine (MDC), macrophage inflammatory protein (MIP)-1α, MIP-1β, MIP-3α, Thymus and activation regulated chemokine (TARC), tumour necrosis factor (TNF)-α, TNF-β, Thymic stromal lymphopoietin (TSLP), vascular endothelial growth factor (VEGF)-A. According to established clinical data treatment(45, 46), any result below the limit of quantification (LOQ) was considered as zero, and if the majority (>75%) of tested OA samples were below the lowest limit of quantification (LOQ), the specific cytokine/chemokine was considered as “non-detectable” and excluded for statistical analysis.

### Selected reaction monitoring LC-MS of lubricin Tn and T antigens

Lubricin (> 90% pure from other glycoproteins) from OA patients (n = 20) were isolated from SF samples by anion exchange chromatography as previously described(47). Lubricin containing fractions were concentrated using 100 kD spinfilters (0.5 mL Amicon Ultra, Millipore) and salt exchange was performed with 3 x 0.5 ml 0.1M NH_4_HCO_3_, followed by vacuum centrifugation to dryness. Lubricin *O*-linked oligosaccharides were released as alditols by reductive β-elimination in 100 μl sodium borohydride (1.0 M) in sodium hydroxide, 0.10 M) at 50°C overnight followed by cleanup on 150 μL cation exchange media (AG50WX8, Bio-Rad, Hercules, CA, US) on top of C18 SPE columns (Strata C18-E, 100mg, Phenomenex, Torrance, CA, US)). The oligosaccharides were dried using vacuum centrifugation, followed by repeated additions of 5 x 50 μL 1% acetic acid in methanol and vacuum centrifugation, to evaporate borate. Glycan standards were reduced to alditols at same conditions as above for 3 hours or overnight.

The oligosaccharides were dissolved in 100 μL MQ-water, followed by injection (2μL) on a Waters UPLC-MS/MS (Acquity Xevo TQ-S) triple quadrupole mass spectrometer. They were separated on porous graphitized carbon columns (100 x 2.1 mm, 3 µm particles, Hypercarb, ThermoFisher Scientific, Waltham, MA, US) kept at 25°C. A standard mixture containing GalNAc-ol (Tn) (Sigma-Aldrich, St Louis, MO, US), and Galβ1-3GalNAc-ol (T) (Dextra, Reading, UK) was prepared by reduction in 1.0 M sodium borohydride. The gradient (36 min) consisted of 0-20 min 0-40% B (A: 10 mM ammonium bicarbonate, B: 80% acetonitrile in 10 mM ammonium bicarbonate), 20-23 min 100% B, 24-26 min wash with 1% acetic acid, then equilibration 26-36 min with 100% A. The flow rate was kept at 150 µL/min. The ESI capillary was kept on 2.5 kV. The source was at 150°C, and the cone at 40V. The instrument was run in positive mode the first 4.35 min for analysis of reduced Tn-antigen (GalNAcol, RT= 4.0 min) and covered the following six transitions at collision energy CE=30 and with a dwell time of 0.052 ms per transition during every cycle: m/z 224 to 182 ([M+H] ^+^-C_2_H_2_O), m/z 224 to 206 ([M+H] ^+^-H_2_0), the C13 isotope transitions: m/z 225 to 183, 225 to 207, and sodium adducts m/z 246 to 182 ([M+Na] ^+^-C_2_H_2_O) and m/z 246 to 206 ([M+Na] ^+^-H_2_O). No sodium adducts were detected. The transition m/z 224 to 182 was chosen for the quantitation calculations applied in this study. During the second period of the chromatographic run the instrument was run in negative mode (4.35-20 min), covering nine different glycan transitions including that for T-antigen (Galβ1-3GalNAc, m/z 384.2 to 204.1 (CE=15) at a dwell time of 0.032 ms. A dilution series (1:2) of the standard mixture was analyzed between every three SF samples, with 0.3-75 pmol injected on the column, using the same method as for the samples. The obtained standard curves for GalNAcol and Galb1-3GalNAcol displayed R^2^ values of 0.9756 (0.5-7.5 pmol) and 0.9948 (2-75 pmol), respectively (Supplementary Figure 2).

### Lubricin-lectin sandwich ELISA

An in-house lectin ELISA was applied for measuring lectin binding on OA synovial fluid lubricin. In brief, the assay buffer was 1% bovine serum albumin (VWR Chemicals, UK) in Tris Buffered Saline (TBS, pH 7.4) with 0.05% tween 20, and the washing buffer is TBS-tween. 96-well plate was coated with 1 μg/ml monoclonal anti-lubricin antibody clone 9G3 (Merck Millipore, Billerica, MA) in TBS at 4°C over-night. After blocking with 3% bovine serum albumin (BSA, VWR Chemicals, UK) in TBS-tween, OA synovial fluid samples that were pre-diluted in assay buffer were loaded to each well in duplicates and incubated for 90 minutes, the captured lubricin was incubated with 1 μg/ml biotinylated Peanut Agglutinin Lectin (PNA) (Vector laboratories, Burlingame, CA, US) or 2 μg/ml biotinylated Helix Aspersa Agglutinin lectin (HAA) (Sigma-Aldrich) followed by 1-hour incubation with 0.2 μg/ml horseradish peroxidase (HRP) conjugated streptavidin (Vector Laboratories). After staining with 1-Step™ Ultra TMB-ELISA Substrate Solution (Thermo Fisher Scientific, San Jose, CA, US), mean absorbances in duplicates were read at 450nm and ratio of HAA/PNA was calculated for each sample in order to measure changes of glycosylation, independent of lubricin level. Recombinant lubricin (rhPRG4, Lubris, LLC, Weston, MA) was used as a standard(48). Validation of the lectin assay showed typically a inter- and intra-plate CV of less than 20% (average 123 μg/mL Lubricin), Linearity of dilution (1/20-1/80) (samples) of more than 70% (2 samples) and spike and recovery of 93% (1.25 μg/mL), 78% (5 μg/mL) and 127% (20 μg/mL) (2 samples) using rhPRG4 for spiking into SF sampels with low lubricin content.

### De-glycosylation of rhPRG4 and treatment of FLS

rhPRG4 was incubated with beta 1-3 galactosidase and alpha 2-3, 6, 8 neuraminidases (both New England BioLabs, Ipswich, MA, US) separately or in combination according to the manufacture’s instructions. FLS(49) were from normal individuals (n=3) and OA patients (n=3) and were incubated with 15 μg of rhPRG4 (intact, de-glycosylated with alpha 2-3, 6, 8 neuraminidases, 1-3 galactosidase or both) for 48 hours at 37 ° C at 5% C0_2_.

### FLS Cytokine expression analysis

FLS were plated (200,000 cells per well) in 6 well dishes 24 hours before the rhPRG4 treatment. FLS were incubated for 48 hours and culture media were collected for cytokine profiling analysis. Cytokine profiling analysis was performed by Eve Technologies (Calgary, AB Canada) using the Milliplex MAP Human Cytokine/Chemokine Panel (Millipore) according to the manufacturer’s instructions. All samples were assayed in duplicate and prepared standards were included in all runs. The following cytokines were quantified in this study: MCP-1, MIP-1β, MIP-1α, IL-8, VEGFA, MDC. The sensitivities of these makers range from 0.1 – 10.1 pg/mL (average 2.359 pg/mL) and the inter-array accuracies ranged from 3.5% – 18.9% coefficient of variation (average 10.7%).

### Statistics

All statistical calculations were performed by GraphPad Prism 8 (GraphPad Software, LLC, USA). In brief, two-tailed non-parametric Spearman correlations was used for calculating correlations. Significant outlier was excluded by ROUT test with Q=0.1%. Each calculation method was specified in the result section. False discovery rate (5%) of individual comparisons was performed for multiple comparisons using Benjamini-Hochberg correction. Significance for secretion of cytokines from FLS was calculated using one way ANOVA multiple comparison.

## Funding

This study was funded by grants for NGK from the Swedish state under the agreement between the Swedish government and the county council, the ALF-agreement (ALFGBG-722391), the Swedish Research Council (621-2013-5895), Petrus and Augusta Hedlund’s foundation (M-2016-0353) and AFA insurance research fund (dnr 150150). GDJ was funded by the National Institute of Health R01AR067748.

## Acknowledgements

Sofia Grindberg and Paula-Therese Kelly Pettersson at Danderyds Hospital and Lotta Falkendahl at University of Gothenburg are acknowledged for their assistance in collecting samples. Lubris BioPharma is acknowledge for the generous gift of rhPRG4.

## Conflict of interest

GJ, RK and TS authored patents related to rhPRG4 and and GDJ and TS hold equity in Lubris BioPharma LLC. TS is also a paid consultant for Lubris BioPharma, LLC. NGK, SH and CJ authored a patent using lubricin and cytokines for diagnostics and NGK and CJ hold equity in Lynxon AB. KAT, HR, AS, ND, OR, LIB and TE declare no conflict of interest.

## Notes

### Summary of Updates

update of Figure 6 and 7 and including extra information in materials and methods

