## Supplementary Figure 1 for "The level of synovial human VEGFA, IL-8 and MIP-1α correlate with truncation of lubricin glycans in osteoarthritis"

**Supplementary Figure 1. Cytokines display different correlation pattern comparing synovial fluid and plasma.** Co-regulation between cytokines concentrations within control plasma (n=16) (A) and plasma, OA plasma (n=29) (B) and OA synovial fluid (n=52) (C). Detectable expressed (above LLOQ) 27 cytokines were correlated within each sample group, and Spearman's rank correlations coefficient r where calculated and indicated in color scale. Positive correlations were in red and negative correlations were in blue as shown in indicator legend. Correlations were calculated by two-tailed Spearman correlation: 1 indicates a perfect positive linear relationship, -1 indicates a perfect negative linear relationship, and 0 stands for no linear relationship. D) Differences in correlation between cytokines were calculated using Spearman's rank correlations coefficients r of the cytokines in each sample were calculated Significances of differences in Spearman's rank between samples were calculated using Mann Whitney U-tests, * indicate p <0.05.

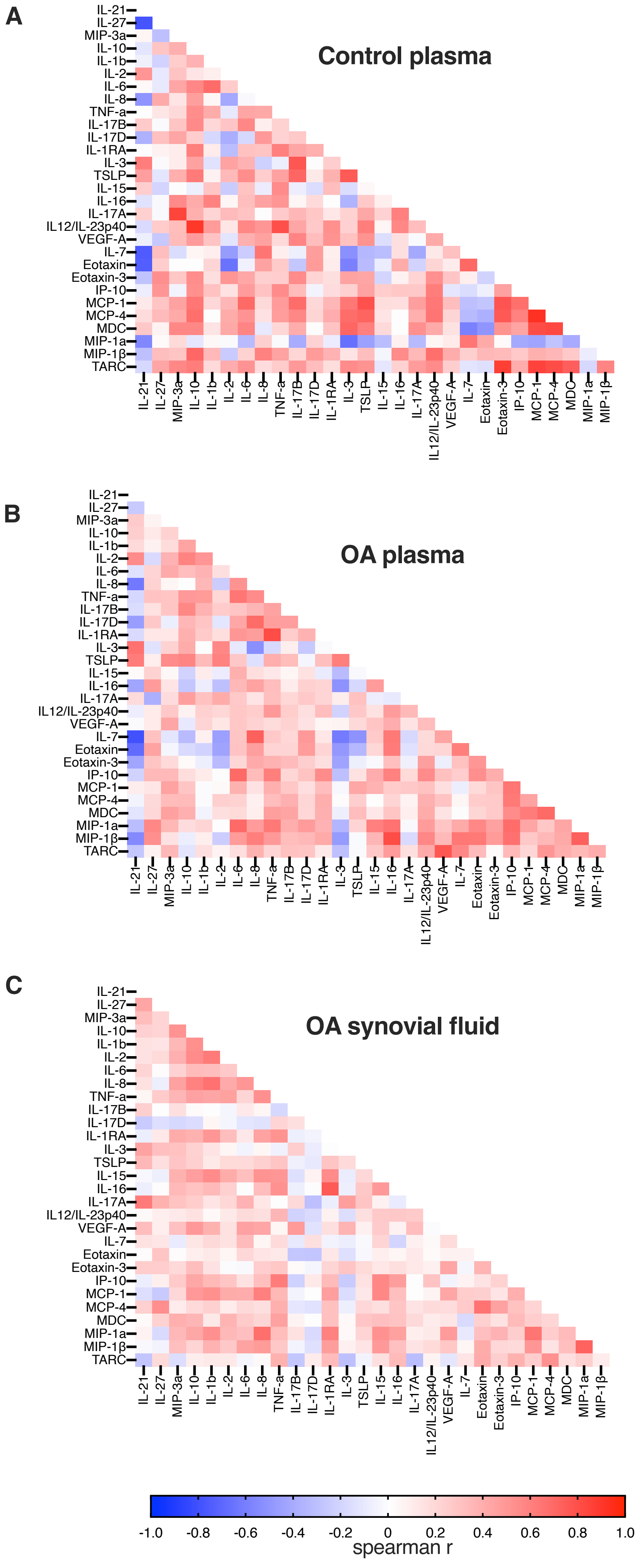


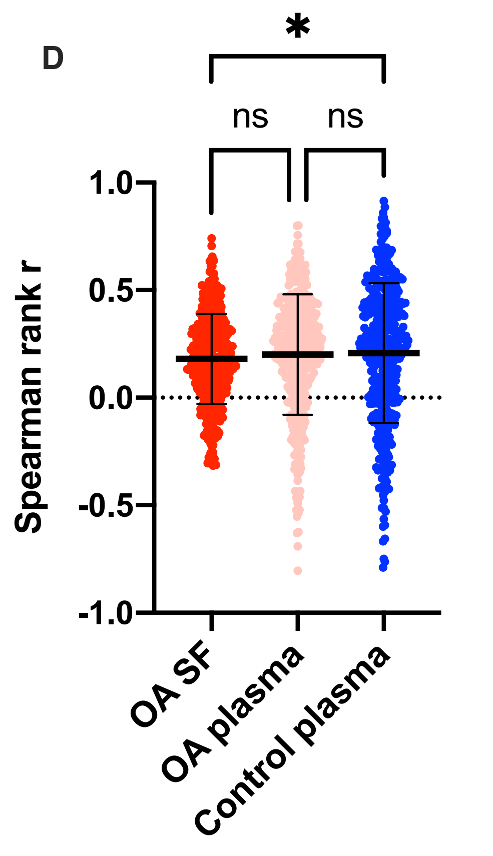
