## Supplementary Figure 2 for "The level of synovial human VEGFA, IL-8 and MIP-1α correlate with truncation of lubricin glycans in osteoarthritis"

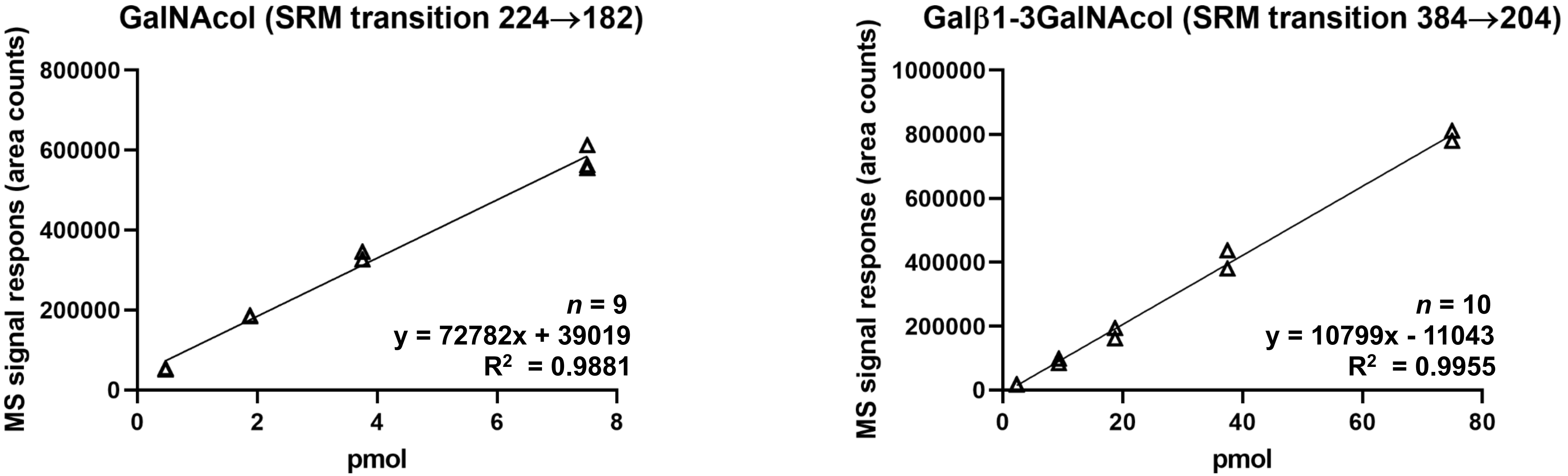


**Supplementary Figure 2**. O-glycans ’GalNAc’ and ’Galβ1-3GalNAc’ were reduced and analyzed as alditols at different concentrations as described in Materials and Methods
