## Supplementary Tables for "The level of synovial human VEGFA, IL-8 and MIP-1α correlate with truncation of lubricin glycans in osteoarthritis"

**Supplementary Table 1.** Inflammatory biomarker concentration differences between OA synovial fluid, OA plasma and control plasma. Differences between control plasma/OA SF and OA plasma/calculatedplasma were calculated by Mann-Whitney nonparametric test with *=p<0.05, **p<0.01, ***p<0.001, ****p<0.0001. Definitive outlier was identified by ROUT with Q=0.1%. Grey boxes indicate values that were shown to be significant after multiple comparison (false discovery rate (5%) using Benjamini-Hochberg correction), comparing values within “OA SF vs OA plasma” and “OA plasma vs control plasma”, respectively.

| | OA SF vs OA plasma | | OA plasma vs control plasma | |
|---|---|---|---|---|
| | Difference between medians (pg/ml) | P value | Difference between medians (pg/ml) | P value |
| Eotaxin | -84.73 | <0.0001 **** | -42.93 | 0.0130 * |
| IP-10 | 14.58 | 0.6155 | 118.3 | 0.0855 |
| MCP-1 | 176.9 | <0.0001 **** | -5.088 | 0.3773 |
| MCP-4 | -52.35 | <0.0001 **** | 26.93 | 0.0010 *** |
| MDC | -624.2 | <0.0001 **** | 204.0 | 0.0497 * |
| MIP-1α | 2.676 | 0.0836 | -2.024 | 0.0426 * |
| MIP-1β | -5.109 | 0.2269 | 3.260 | 0.4648 |
| TARC | -84.01 | <0.0001 **** | 68.71 | 0.0030 ** |
| IL-15 | 14.45 | <0.0001 **** | -0.2388 | 0.7559 |
| IL-16 | 85.62 | 0.0002 *** | 19.68 | 0.9906 |
| IL-7 | -1.158 | 0.2442 | 2.605 | 0.4193 |
| IL-12/IL-23p40 | -65.88 | <0.0001 **** | -69.50 | 0.1551 |
| VEGF-A | 353.0 | <0.0001 **** | 0.8323 | 0.8420 |
| IL-17B | 32.16 | <0.0001 **** | 0.1719 | 0.4147 |
| IL-17D | 97.17 | <0.0001 **** | 1.026 | 0.4788 |
| IL-1RA | -44.41 | 0.1856 | 64.13 | 0.0417 * |
| IL-3 | -9.672 | 0.0093 ** | 0.9920 | 0.9347 |
| TSLP | 0.04972 | 0.8677 | -0.02534 | 0.5069 |
| IL-10 | -0.03271 | 0.2383 | -0.04859 | 0.9473 |
| IL-6 | 22.44 | <0.0001 **** | 0.2738 | 0.0688 |
| IL-8 | 20.16 | <0.0001 **** | 1.350 | 0.0179 * |
| TNF-α | -1.448 | <0.0001 **** | 0.3595 | 0.3059 |
| IL-21 | -5.876 | <0.0001 ****[a] | -1.139 | 0.6953 |
| IL-27 | -1259 | <0.0001 **** | 126.1 | 0.3529 |
| MIP-3α | -1.152 | 0.0395 * | -0.3265 | 0.7295 |
| IL-1β | 0.07391 | 0.1541 | 0.02970 | 0.5452 |
| IL-1α | -2.292 | <0.0001 ****[a] | -0.3974 | 0.4830 |
| IL-2 | 0.2529 | <0.0001 **** | -0.05807 | 0.9248 |
| Eotaxin-3 | -12.94 | <0.0001 **** | 3.796 | 0.0267 * |

[a]Measurement was generally low in SF of these cytokines and was excluded applying our exclusion criteria, but was included here because of consistency of the table, and to illustrate that these cytokines were very low in SF compared to plasma

**Supplementary Table 2.** Cytokine expressions in OA and control SF

| Cytokine | OA (pg/ml) N=52 | Control (pg/ml) N=3 | OA Literature (pg/ml) | Control literature (pg/ml) |
|---|---|---|---|---|
| IL-17A Gen.B | 2.475 ± 2.794 | 2.704 ± 4.683 | | |
| IL-21 | 1.61 ± 2.405 | 1.345 ± 2.33 | | |
| IL-27 | 92.47 ± 52.83 | 100.5 ± 76.8 | | |
| MIP-3a | 6 ± 3.152 | 530 ± 854.8 | | |
| IL-10 | 0.6197 ± 1.782 | 1.535 ± 1.399 | 9.1 ± 10.5[1], 9 ± 35[2] | 1 ± 6[2] |
| IL-1β | 0.7774 ± 1.71 | 11.92 ± 17.78 | 8 ± 16[2] | 1 ± 2[2] |
| IL-2 | 0.8598 ± 0.3334 | 4.481 ± 5.164 | | |
| IL-6 | 42.79 ± 55.92 | 22.66 ± 39.03 | 8.3 ± 11.1[1], 396 ± 508[2], 53[3] | 64 ± 120[2] |
| IL-8 | 64.49 ± 96.5 | 21.54 ± 31.09 | 8.2 ± 14.5[1], 52 ± 95[2], 36[3] | 25 ± 29[2] |
| TNF-a | 0.9998 ± 0.4758 | 3.658 ± 3.46 | 4 ± 20[2] | 0 ± 0[2] |
| IL-17B | 45.17 ± 37.05 | 39.92 ± 26.67 | | |
| IL-17D | 165.6 ± 227.5 | 586.3 ± 463.4 | | |
| IL-1RA | 323.8 ± 342.6 | 66.13 ± 104.56 | 560[3] | |
| IL-3 | 26.38 ± 63.03 | 20.29 ± 7.03 | | |
| TSLP | 1.5 ± 1.229 | 6.075 ± 8.05 | 5.17[4] | 5.05[4] |
| IL-15 | 17.99 ± 4.931 | 16.91 ± 5.342 | | |
| IL-16 | 533.7 ± 585.6 | 233.2 ± 257.1 | | |
| IL-17A | 1.988 ± 1.389 | 1.923 ± 0.7102 | | |
| IL-7 | 5.385 ± 2.274 | 2.663 ± 1.59 | 5 ± 28[2],1.3 – 2.6[3] | 0 ± 0[2] |
| IL-12 | 88.11 ± 49.76 | 37.53 ± 19.58 | | |
| VEGF-A | 515.2 ± 370.2 | 575.3 ± 544.7 | 490 – 1000[3] | |
| Eotaxin | 39.24 ± 14.17 | 51.66 ± 12.85 | 1.5 ± 1.0[1] | |
| Eotaxin-3 | 3.548 ± 3.954 | 8.947 ± 6.032 | | |
| IP-10 | 977.9 ± 249.2 | 1814.2 ± 2731.6 | | |
| MCP-1 | 347.5 ± 187.8 | 318.6 ± 151.2 | 30.3 ± 18.9[1], 300[3] | |
| MCP-4 | 18.89 ± 18.69 | 227.2 ± 151.4 | | |
| MDC | 391.3 ± 109.3 | 26.43 ± 14.07 | | |
| MIP-1a | 17.98 ± 10.69 | 163.7 ± 57.37 | 34[3], 700 ± 400[5] | |
| MIP-1β | 47.22 ± 22.06 | 15.13 ± 4.334 | | |
| TARC | 21.01 ± 16.08 | 13.38 ± 2.08 | 1.4 ± 0.5[1], >10[6] | |
| IL-22 | 0.8085 ± 0.6876 | 22.05 ± 8.378 | | |
| IL-23 | 0 ± 0 | 1.908 ± 0.487 | | |
| IL-31 | 0.2871 ± 0.4122 | 15.71 ± 15.86 | | |
| IFN-γ | 1.696 ± 4.682 | 0.4463 ± 0.6125 | | |
| IL-12p70 | 0.1581 ± 0.1885 | 21.27 ± 31.61 | | |
| IL-13 | 1.289 ± 1.029 | 14.42 ± 23.34 | | 1 ± 2[2] |
| IL-4 | 0.1041 ± 0.1025 | 16.75 ± 10.79 | | 0 ± 0[2] |
| IL-17A/F | 0.4217 ± 0.8815 | 6.119 ± 9.759 | | |
| IL-17C | 3.881 ± 20.27 | 1.816 ± 1.905 | | |
| IL-9 | 0.3264 ± 0.2339 | 5.68 ± 4.389 | | |
| GM-CSF | 0.2232 ± 0.1994 | 0.802 ± 0.5568 | | |
| IL-1α | 0.1146 ± 0.5449 | 0.5647 ± 0.2758 | 15 ± 22[2] | 16 ± 10[2] |
| IL-5 | 0.1313 ± 0.2068 | 1.06 ± 0.4126 | | |
| TNF-β | 0.2546 ± 0.2588 | 0.488 ± 0.4541 | | |

Concentrations presented as mean± SD, non-detectable listed as 0.

**Supplementary Table 3.** Correlation between plasma and SF expressed inflammatory biomarkers in 28 OA patients.

| Inflammatory biomarker | Spearman r | P value |
|---|---|---|
| *Th17 pathway cytokines* | | |
| IL-27 | 0.3893 | 0.0203 * |
| IL-21 | 0.3131 | 0.0729 |
| MIP-3a | 0.1671 | 0.2073 |
| *Proinflammatory cytokines and chemokines* | | |
| IL-12 | 0.5863 | 0.0008 *** |
| IL-10 | 0.4607 | 0.0089 ** |
| TNF-a | 0.6268 | 0.0002 ** |
| IP-10 | 0.6504 | 0.0003 *** |
| MDC | 0.5025 | 0.0032 ** |
| IL-1b | -0.03219 | 0.8869 |
| IL-2 | -0.2970 | 0.0703 |
| IL-6 | 0.06609 | 0.7590 |
| IL-15 | 0.2151 | 0.1358 |
| IL-16 | 0.2479 | 0.1111 |
| VEGF-A | -0.2155 | 0.1402 |
| IL-8 | 0.1070 | 0.3094 |
| Eotaxin | 0.08518 | 0.3364 |
| MCP-1 | 0.04783 | 0.4122 |
| MCP-4 | 0.3128 | 0.0599 |
| MIP-1a | 0.1870 | 0.1854 |
| TSLP | 0.4032 | 0.0205 * |
| IL-17B | -0.08487 | 0.3369 |
| IL-17D | -0.04040 | 0.4240 |
| IL-1RA | 0.2348 | 0.1145 |
| IL-3 | 0.3998 | 0.0501 |
| IL-7 | 0.1520 | 0.2245 |
| MIP-1b | 0.2950 | 0.0637 |
| TARC | 0.2308 | 0.1283 |

P value and corresponding significance for the correlation are shown according to nonparametric Spearman correlation with *=p<0.05, **p<0.01, ***p<0.001, ****p<0.0001. Grey boxes symbolizes values that were shown to be significant after multiple comparison (false discovery rate (5%) using Benjamini-Hochberg correction).

**Supplementary Table 4.** Correlation between HAA/PNA ratio and SF expressed inflammatory biomarkers in 28 OA patients.

| Inflammatory biomarker | Spearman r | P value |
|---|---|---|
| IL-21 | 0.1645 | 0.4123 |
| IL-27 | -0.02057 | 0.9125 |
| MIP-3α | 0.1517 | 0.4154 |
| IL-10 | 0.2065 | 0.2918 |
| IL-1b | 0.173 | 0.3788 |
| IL-2 | -0.1688 | 0.3813 |
| IL-6 | 0.1445 | 0.4632 |
| IL-8 | 0.5244 | 0.0042** |
| TNF-α | -0.07313 | 0.7115 |
| IL-17B | 0.1858 | 0.3257 |
| IL-17D | -0.3637 | 0.0482* |
| IL-1RA | 0.1528 | 0.4201 |
| IL-3 | 0.2507 | 0.2072 |
| TSLP | 0.2072 | 0.2807 |
| IL-15 | 0.204 | 0.2709 |
| IL-16 | -0.03938 | 0.8363 |
| IL-7 | 0.07298 | 0.6964 |
| IL12/IL-23p40 | 0.1346 | 0.4783 |
| VEGF-A | 0.7433 | <0.0001**** |
| Eotaxin | -0.03871 | 0.8362 |
| IP-10 | 0 | >0.9999 |
| MCP-1 | 0.4202 | 0.0186* |
| MCP-4 | -0.017 | 0.9303 |
| MDC | 0.3738 | 0.0383* |
| MIP-1α | 0.6114 | 0.0003*** |
| MIP-1β | 0.4136 | 0.0231* |
| TARC | -0.08677 | 0.6484 |

P value and corresponding significance for the correlation are shown according to nonparametric Spearman correlation with *=p<0.05, **p<0.01, ***p<0.001, ****p<0.0001. Grey boxes shows values that were shown to be significant after multiple comparison (false discovery rate (5%) using Benjamini-Hochberg correction).
